# Hemangiosarcoma cells induce M2 polarization and PD-L1 expression in macrophages

**DOI:** 10.1101/2021.08.01.454679

**Authors:** Kevin Christian M. Gulay, Keisuke Aoshima, Naoya Maekawa, Satoru Konnai, Atsushi Kobayashi, Takashi Kimura

## Abstract

Hemangiosarcoma (HSA) is a malignant tumor derived from endothelial cells. Tumor-associated macrophages are one of the major components of tumor microenvironment and crucial for cancer development. The presence and function of macrophages in HSA have not been studied because there is no syngeneic model for HSA. In this study, we evaluated two mouse HSA cell lines and one immortalized mouse endothelial cells for their usefulness as syngeneic models for canine HSA. Our results show that the ISOS-1 cell line develops tumors with similar morphology to canine HSA. ISOS-1 cells highly express KDM2B and have similar KDM2B target expression patterns with canine HSA. Moreover, we determine that in both ISOS-1 and canine HSA tumors, macrophages are present as a major constituent of the tumor microenvironment. These macrophages are positive for CD204, an M2 macrophage marker, and express PD-L1. ISOS-1-conditioned medium can induce M2 polarization and PD-L1 expression in RAW264.7 mouse macrophage cell line. These results show that ISOS-1 can be used as a syngenic model for canine HSA and suggest that macrophages play an important role in immune evasion in HSA. Using the syngeneic mouse model for canine HSA, we can further study the role of immune cells in the pathology of HSA.

## Introduction

Hemangiosarcoma (HSA) is a rapidly growing and highly invasive endothelial cancer^1^. It is the most common splenic neoplasm in dogs where it usually develops at 6 to 17 years of age^2^. Middle to large breed dogs are most commonly afflicted with HSA^2^. HSA also occurs, albeit infrequently, in cats, horses, mice, and humans^3–6^. An effective treatment for HSA is difficult to develop since little is known about its molecular pathology. Recently, we found that canine HSA highly expressed three histone demethylases (KDM1A, KDM2A and KDM2B) out of which KDM2B was found necessary for HSA cell survival by positively regulating the DNA damage response system in tumor cells^7^. KDM2B silencing not only dysregulated DNA damage response but also induced expressions of the genes related to inflammatory responses^7^. Inhibiting KDM2B could be an option to induce host immune responses against HSA tumor cells. However, we were unable to investigate the function of KDM2B in canine HSA tumor immune responses because the immunodeficient mouse model was not suitable to study the immune responses^8^. Syngeneic mouse models, otherwise known as allograft mouse tumor models, are composed of tumor tissues derived from the same genetic background as the mouse strain. The syngeneic mouse models can develop tumors in a fully immunocompetent environment, which can facilitate the examination of the immune-tumor cell interactions. A syngeneic model for HSA, however, is non-existent.

At present, there are few established mouse HSA cell lines such as ISOS-1 and UV♀2. ISOS-1 is a mouse HSA cell line established from a tumor formed by the xenotransplantation of a human angiosarcoma cell line, while UV♀2 cell line is a mouse HSA cell line developed from an ultraviolet light-induced HSA^9,10^. Their usefulness as a syngeneic model for canine HSA or human angiosarcoma, however, has not been evaluated.

Macrophages have two states, M1 and M2. M1 macrophages induced by Lipopolysaccharides (LPS) and IFNγ are capable of killing tumor cells and presenting tumor antigens to CD4^+^ T cells^11^. M2 macrophages are polarized by IL-4, IL-10 and TGFβ and are important for tumor growth and immune evasion^11^. They can produce anti-inflammatory cytokines like IL-10, IL-3, and TGF-β which supports tumor development by repressing cytotoxic T cell function^12,13^. M2 macrophages in tumor tissues can be detected using CD204 (also known as MSR1) as a M2 macrophage marker^14^. *In vitro*, CD204 expression is induced by TGFβ treatment but LPS-treatment can also induce its expression via the MAPK/ERK pathway in murine bone marrow-derived macrophages^15,16^. It is highly likely that M2 macrophages support tumor development and facilitate immune evasion in HSA; however, there are no studies that have evaluated their presence and functions in HSA.

In this study, we aimed to evaluate existing mouse HSA cell lines and immortalized endothelial cells for their possible use as a syngeneic model for canine HSA and to identify the constituents of the tumor microenvironment and their roles in canine and mouse HSA pathology.

## Results

### ISOS-1 cells can be used as a syngenic model of canine HSA

To find syngeneic models for canine HSA, we first characterized two mouse endothelial tumor cell lines: ISOS-1 and UV♀2, and one immortalized mouse endothelial cell line, LEII. Morphologically, in single layer culture, both the mouse and canine HSA cell lines are characterized by spindle to polygonal cells with a moderate amount of cytoplasm and elongate nuclei arranged in cobblestone pattern (Fig. S1A). Then, we inoculated ISOS-1, LEII and UV♀2 into Balb/c mice subcutaneously to determine their tumorigenic potential. All mice inoculated with ISOS-1 cells developed tumors after 30 days post inoculation (dpi) and reached the endpoint before 65 dpi (Fig. S1B). Mice inoculated with UV♀2 or LEII cells, however, did not develop tumors. No metastasis was observed in ISOS-1 inoculated mice at the endpoint. We then made histopathological sections of ISOS-1 tumors and compared their morphologies with canine clinical HSA cases. ISOS-1 tumor cells formed variably sized, irregular shaped blood vessels that were separated by thin septa and trabeculae. They showed solid and capillary growth patterns which were also observed in canine HSA (Fig. 1A). Spindle-shaped neoplastic endothelial cells line luminal spaces in a single layer and are plump, hyperchromatic, and larger than normal endothelial cells (Fig. 1A, insets).

**Fig. 1.**
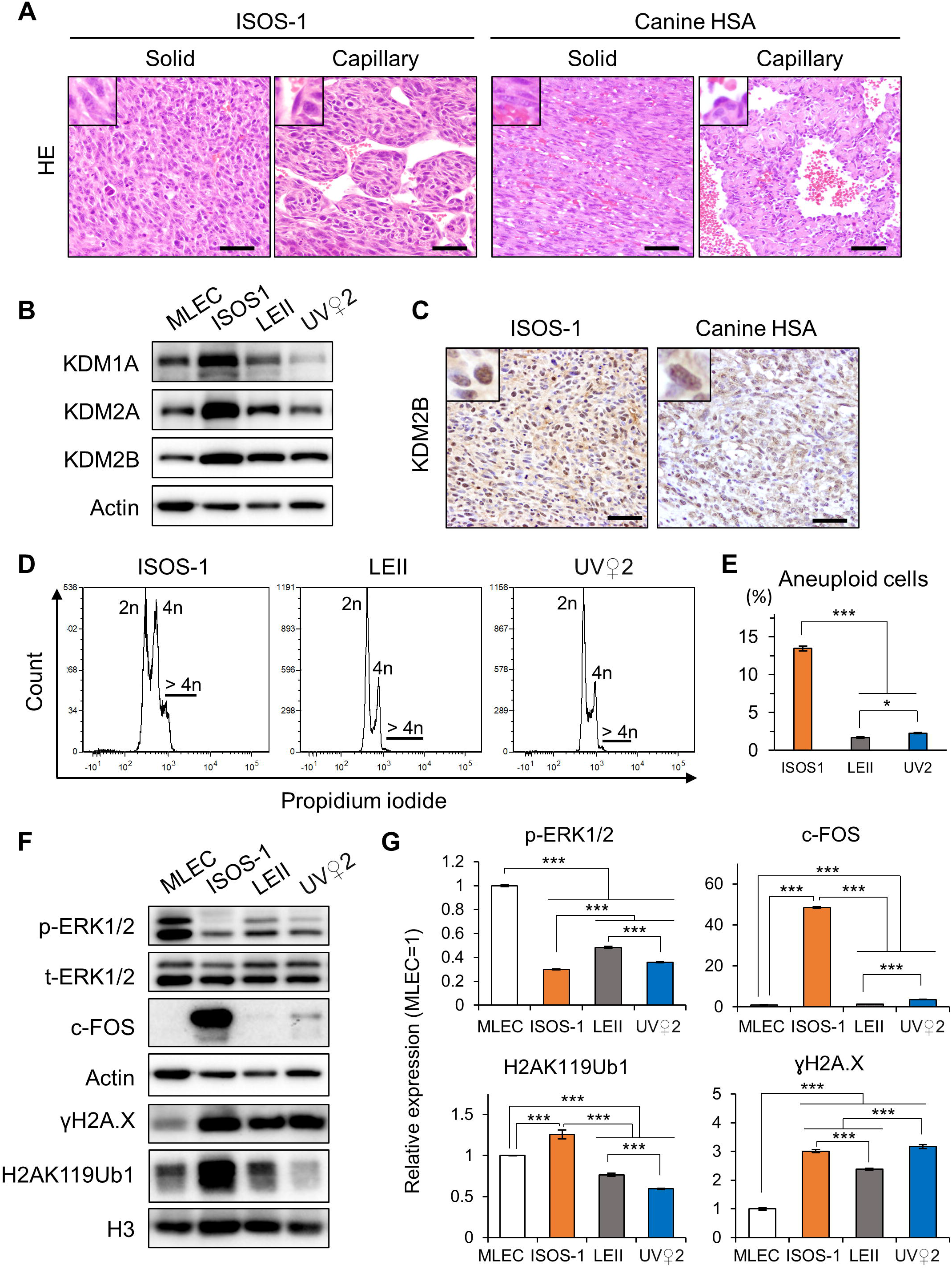
ISOS-1 presents similar morphology and molecular features with canine HSA. **A**, Hematoxylin and eosin (HE) staining in ISOS-1 and canine HSA tumors. **B**, KDM1A, KDM2A, and KDM2B protein expressions in MLEC, ISOS1, LEII, and UV♀2 cell lines. **C**, KDM2B immunohistochemistry of ISOS-1 and canine HSA tumors. **D**, Histograms of PI intensities in ISOS1, LEII, and UV♀2 cell lines. **E**, Percentages of cells with aneuploidy in ISOS1, LEII, and UV♀2 cell lines. **F**, Western blotting for phosphorylated ERK1/2(p-ERK1/2), c-FOS, γH2A.X and H2AK119ub1 in MLEC, ISOS-1, LEII, and UV♀2 cell lines. **G**, Quantification of p-ERK1/2, c-FOS, γH2A.X, and H2AK119ub1 in MLEC, ISOS-1, LEII, and UV♀2 cell lines. p-ERK1/2 expression was normalized with total ERK1/2 expression. c-FOS expression was normalized with Actin expression. H2AK119Ub1 and γH2A.X expressions were normalized with H3 expression levels. The protein expression levels in MLEC were set to 1. Data are presented as mean values ± s.d. Experiments were performed in triplicates. Scale = 125 μm. ^***^ *P* < 0.001, Tukey’s test.

Next, we investigated whether ISOS-1, LEII and UV♀2 have similar molecular characteristics with canine HSA. Although *Kdm1a, Kdm2a* and *Kdm2b* gene expressions in the mouse endothelial cell lines were not expressed more than 2-fold compared with primary mouse lung endothelial cells (MLEC), their protein expressions in ISOS-1 were significantly higher than in MLEC, LEII and UV♀2 (Fig 1B and Fig. S2A). ISOS-1 tumor cells in Balb/c mice also expressed KDM2B as high as in canine HSA (Fig 1C). Furthermore, ISOS-1 cells had other similar molecular features to canine HSA cell lines such as cellular aneuploidy, low p-ERK expression level and high c-FOS, γH2A.X and H2AK119Ub1 expression levels (Figs. 1D - 1G).

Finally, we treated the mouse endothelial cell lines with GSK-J4, a histone demethylase inhibitor, to know whether GSK-J4 can inhibit their viability and whether it is more effective than doxorubicin like in canine HSA. The results showed that GSK-J4 could inhibit the cell viability of ISOS-1, LEII and UV♀2 at a lower IC_50_ compared to doxorubicin (Fig. S2).

These results demonstrate the numerous similarities between ISOS-1 cell line and canine HSA, and they provide more proof of the usefulness of ISOS-1 as a syngeneic model for canine HSA.

### CD204^+^ macrophage is the major constituent in HSA tumor microenvironment

To identify the constituent cells of the tumor microenvironment in HSA, we immunohistochemically stained four ISOS-1 tumors developed in Balb/c mice and twenty-eight clinical HSA samples in dogs with antibodies for Iba1 and CD3, a macrophage and a T cell marker, respectively. T cells were present in minimal amount and were mostly confined in the peripheries of tumor tissues. However, Iba1 staining revealed that macrophages were the major components of tumor microenvironment in both ISOS-1 tumor and canine HSA cases (Fig. 2A). The average percentages of macrophages in tumor tissues were 65.27% and 48.83% in ISOS-1 tumors and canine HSA cases, respectively (Figs. 2B).

**Fig. 2.**
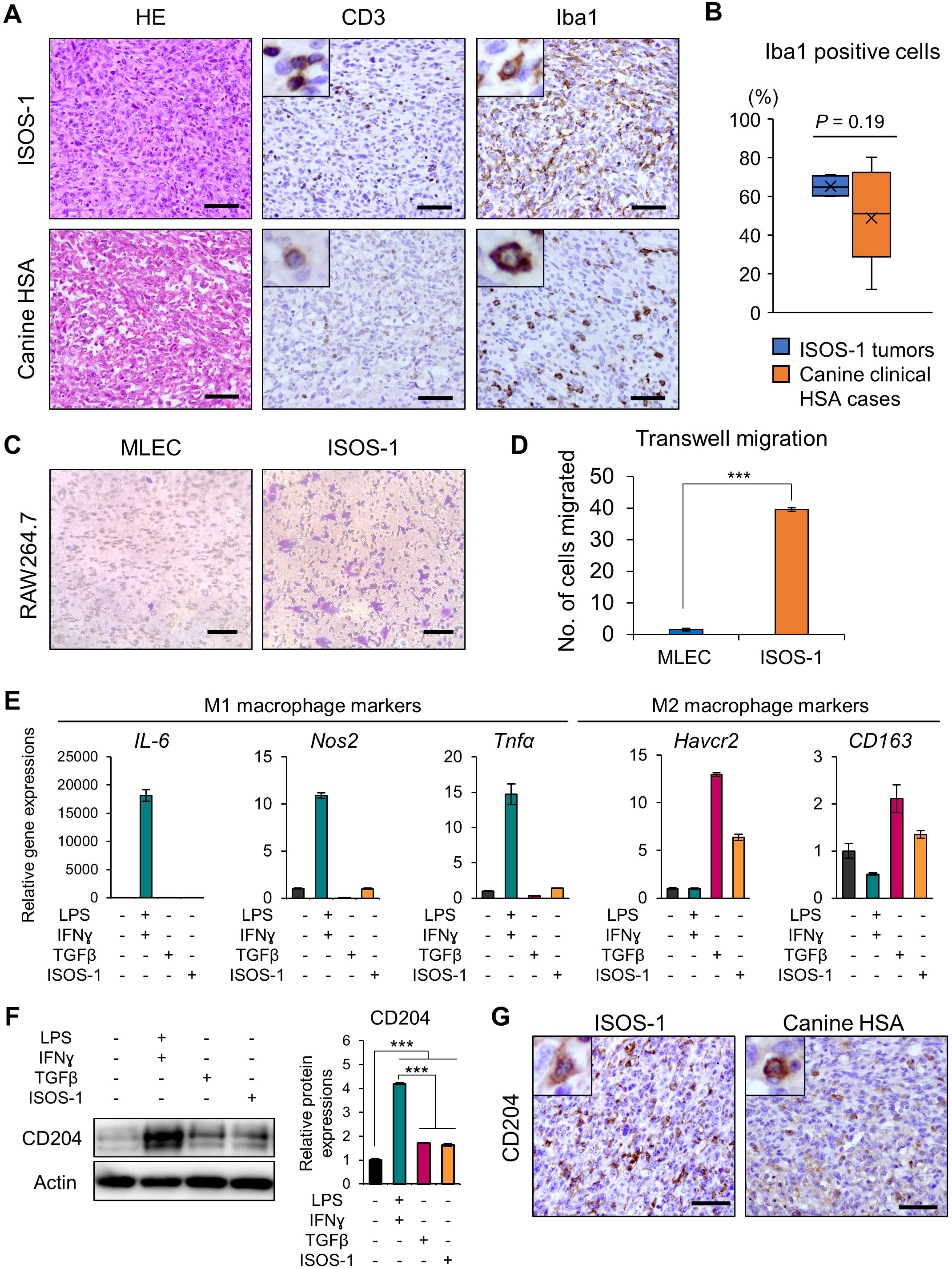
HSA cells attract macrophages and polarize them to M2 macrophages. **A**, HE staining and immunohistochemistry of CD3 and Iba1 for ISOS-1 and canine HSA tumors. **B**, Quantitative analysis of Iba1 positive cells in ISOS-1 and canine HSA tumors. Y-axis indicates the percentages of Iba1 positive cells relative to all cells comprising the tumor tissue. **C**, Transwell migration assay in RAW264.7 cells cultured over MLEC-or ISOS-1-conditioned media. **D**, Quantitative analysis of C. **E**, Gene expressions of M1 or M2 macrophage markers in untreated, LPS and IFN-treated, TGFβ-treated, and ISOS-1 conditioned media-treated RAW264.7 cells. **F**, (Left) Western blot analysis for CD204 in untreated, LPS and IFN-treated, TGFβ-treated, and ISOS1 conditioned media-treated RAW264.7 cells. (Right) Quantitative analysis of the western blotting data. **G**, Immunohistochemistry of CD204 for ISOS-1 and canine HSA tumors. Scale = 125 μm. Data are presented as mean values ± s.d.

Next, to know whether HSA cells actively recruit macrophages, we performed Transwell migration assay and compared the number of migrated macrophage-like RAW264.7 cells in conditioned media from MLEC or ISOS-1. As a result, the number of migrated RAW264.7 cells significantly increased when cultured in conditioned medium from ISOS-1 compared to the conditioned medium from MLEC (Figs. 2C and 2D). RAW264.7 cells cultured in ISOS-1 conditioned medium did not induce M1 macrophage-related genes but highly expressed M2 macrophage markers such as *HAVCR2* and *CD163* (Fig. 2E). The protein expression of CD204, another M2 macrophage marker, was also induced by ISOS-1 conditioned medium in RAW264.7 cells. Lastly, we stained ISOS-1 tumors and canine HSA cases with anti-CD204 antibody and identified a large number of CD204 positive cells in ISOS-1 tumors and canine HSA samples (Fig. 2G)

These results suggest that HSA tumor cells recruit macrophages into the tumor microenvironment and polarize them to M2 macrophages.

### Tumor cells and tumor infiltrating macrophages express PD-L1 in HSA

In both ISOS-1 tumor and canine HSA cases, macrophages dominate the tumor parenchyma along with the tumor cells, thus, it is vital to know how these macrophages contribute to HSA pathology. CD204^+^ macrophages in tumor tissues have been reported to express PD-L1 and are associated with tumor malignancy and PD-L1 upregulation in tumor and immune cells^17,18^. Thus, we decided to examine PD-L1 expressions in tumor cells and macrophages in clinical HSA cases and HSA cell lines.

PD-L1 was expressed in tumor cells and macrophages in 15 out of the 28 (53.6%) and 19 out of the 28 (67.8%) canine HSA cases, respectively (Fig. 3A and Table). 23 out of 28 cases (82.1%) expressed PD-L1 in tumor cells and/or macrophages (Table). The PD-L1 siganl intensity in macrophages was stronger than that in ISOS-1 tumor cells. PD-L1 was also expressed in ISOS-1 and canine HSA cell lines: JuB2, JuB4 and Re21 (Figs 3B and 3C, Figs S3A and S3B). The number of PD-L1 positive cells was slightly increased by IFNγ treatment in ISOS-1 whereas almost 100% of canine HSA cells expressed PD-L1 without IFNγ treatment (Figs. 3B and C; Figs. S3A and 3B). Double staining of PD-L1 and Iba1 verified that macrophages in ISOS-1 tumors and canine HSA cases expressed PD-L1 (Fig. 4A). Then, we tested whether HSA tumor cells can induce PD-L1 expression in macrophages using RAW264.7 cells. PD-L1 gene expression in RAW264.7 cells was induced by LPS and IFNγ treatment (Fig. 4B). Furthermore, ISOS-1 conditioned medium significantly induced PD-L1 protein expression in RAW264.7 cells (Figs. 4C and 4D).

**Fig. 3.**
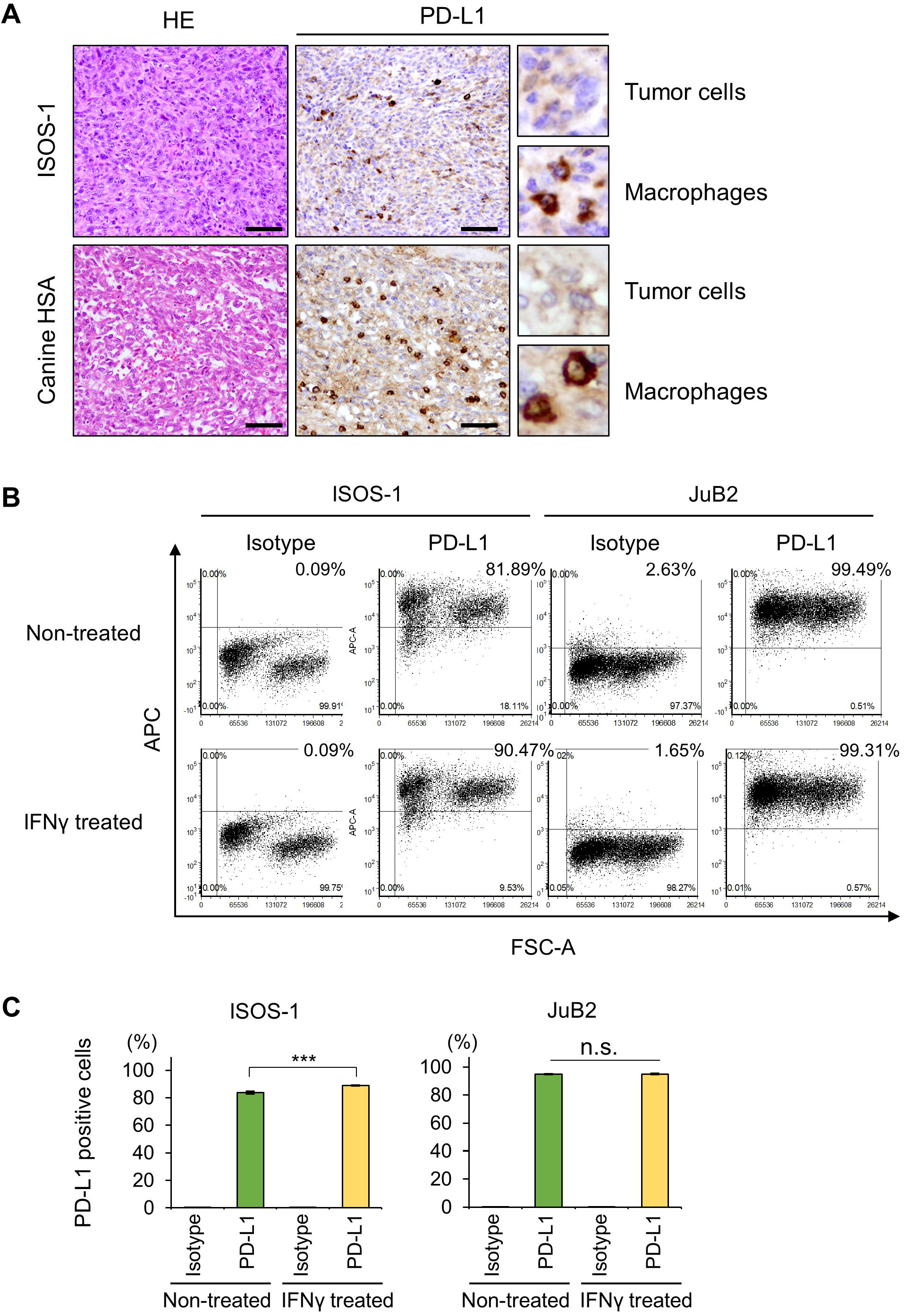
HSA cells express PD-L1. **A**, HE staining and immunohistochemistry of PD-L1 for ISOS-1 and canine HSA tumors. HE staining images are the same as in Fig. 2A because the same samples were used for this experiment. **B**, Representative images of flow cytometry analysis for PD-L1 in ISOS-1 and JuB2 cell lines with/without IFNγ treatment. APC indicates PD-L1 expressions. **C**, Quantitative analysis of B. Scale = 125 μm. Data are presented as mean values ± s.d. ^***^ *P* < 0.001, Tukey’s test.

**Fig. 4.**
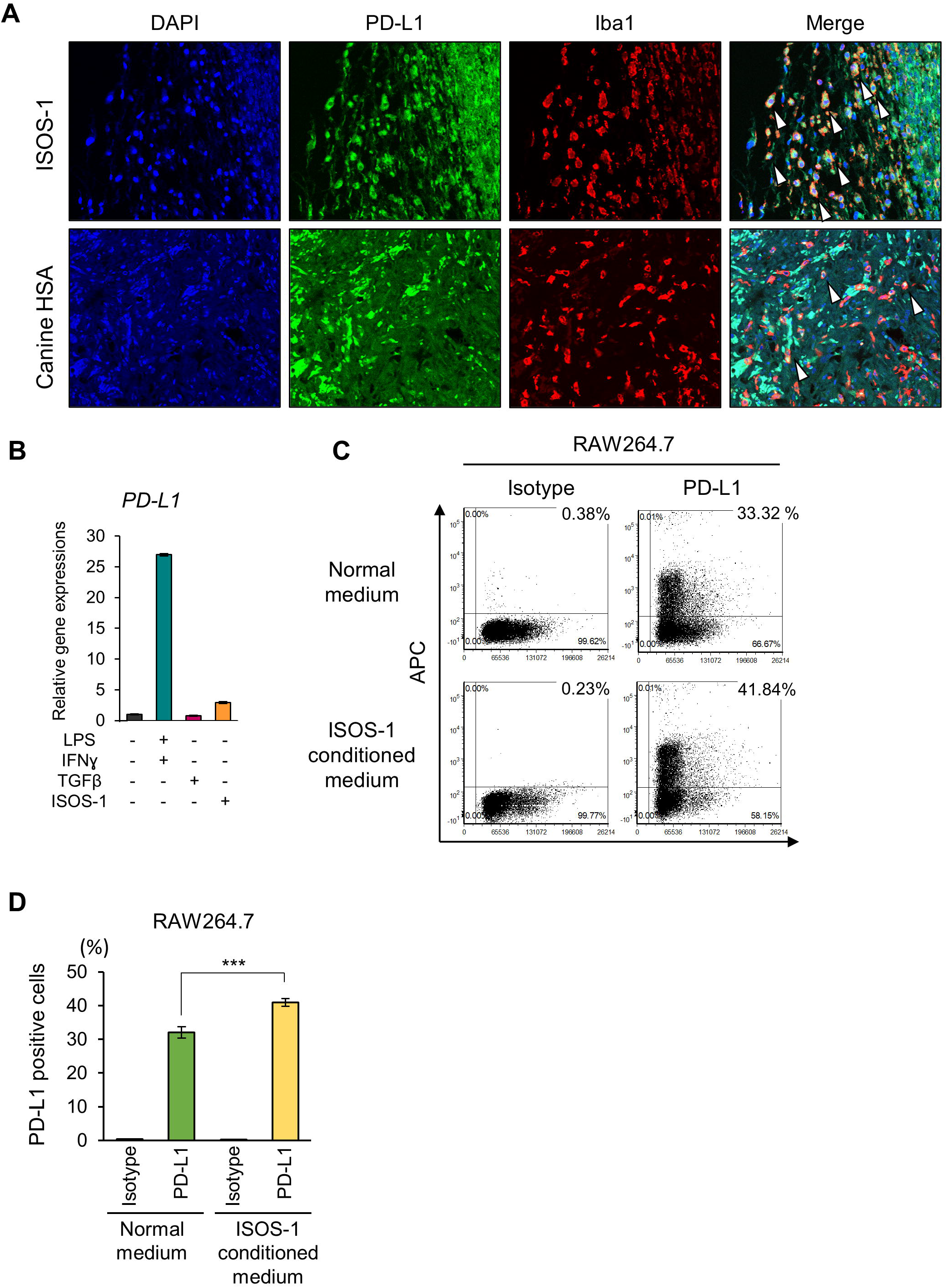
ISOS-1 cells induce PD-L1 expression in RAW264.7 cells. **A**, Immunofluorescence assay for PD-L1 and Iba1 in ISOS-1 and canine HSA tumors. Arrows indicate the cells expressing both PD-L1 and Iba1. **B**, Gene expression levels of PD-L1 in untreated, LPS and IFN-treated, TGFβ-treated, and ISOS-1 conditioned media-treated RAW264.7 cells. **C**, Representative images of flow cytometry analysis for PD-L1 in RAW264.7 cells cultured in normal or ISOS-1 conditioned media. **D**, Quantitative analysis of C. Data are presented as mean values ± s.d. ^***^ *P* < 0.001, Tukey’s test.

These results suggest that HSA evade immune attack through PD-L1 expression in tumor cells and by inducing PD-L1 expression in macrophages.

## Discussion

Here we demonstrated that a mouse HSA cell line, ISOS-1, can be used as a syngenic model for HSA, and that macrophages are the major constituent of the HSA tumor microenvironment in both ISOS-1 and canine HSA tumors. We also identified that ISOS-1 cells could recruit macrophages, polarize them to M2 macrophages, and induce PD-L1 expression in macrophages. In this study, we used two mouse HSA cell lines (ISOS-1 and UV♀2) and one immortalized mouse endothelial cell line (LEII) as candidates of syngenic models for canine HSA. As previous studies reported, ISOS-1 was the only mouse HSA cell line that developed tumors in immunocompetent mice ^9,19^. ISOS-1 possessed similar molecular features with canine HSA such as high KDM2B expression and similar expression patterns of KDM2B targets whereas KDM2B was not highly expressed in UV♀2 and LEII. Based on these results and our previous findings that KDM2B plays an important in canine HSA, it is highly likely that KDM2B is a common factor for endothelial cell tumor malignancy.

We demonstrated that ISOS-1 cells recruited macrophages, polarized them to M2 macrophages, and induced PD-L1 expression in macrophages. In canine HSA tumors, macrophages expressed both CD204^+^ and PD-L1, which suggests that canine HSA cells also attract and induce macrophages to express PD-L1. Since T cells were located at the periphery of canine HSA cases, macrophages in canine HSA likely facilitate immune evasion by through induction of PD-L1 expression. Antibodies specific for canine PD-L1 have been developed and have been tested for their safety and efficacy in clinical cases^20–22^. However, anti-PD-L1 antibody treatment has not been studied in canine HSA patients. Given that more than 80% of clinical HSA cases that we examined in our study expressed PD-L1 in tumor cells and/or macrophages, immunotherapy using anti PD-L1 antibody treatment could be useful as an alternative treatment for canine HSA. In our previous study, silencing of KDM2B resulted to increased interferon gamma and alpha responses^7^. This means that KDM2B inhibition induces immune reaction; therefore, combination therapy with anti-PD-L1 antibody and KDM2B inhibition might provide better outcomes than single treatment with anti-PD-L1 or KDM2B inhibitor treatment in canine HSA.

In summary, we identified the similarities between ISOS-1 and canine HSA, and we demonstrated the usefulness of ISOS-1 as a syngeneic model for canine HSA. By taking advantage of ISOS-1 cells, we characterized the tumor microenvironment in HSA and demonstrated the crosstalk between tumor cells and macrophages for the induction of PD-L1 expression. These results provide useful insights for understanding HSA pathology and will be beneficial to develop novel therapeutics for HSA.

## Materials and Methods

### Cell lines

ISOS-1 cells were obtained from the Cell Resource Center for Biomedical Research Cell Bank (Tohoku University)^9^. UV♀2 cells were obtained from RIKEN Bioresource Center^10^. The LEII cell line was donated by Dr. Kazuhiro Kimura (Hokkaido University) and cultured as described previously^23, 24^. RAW264.7 cells were obtained from RIKEN Bioresource Center^25^. Canine HSA cell lines (JuB2, JuB4, Re12, Ud6) were given by Dr. Hiroki Sakai (Gifu University)^26^. All cells used were routinely tested for *Mycoplasma* using PCR and were submitted to ICLAS Monitoring Center (Kawasaki, Japan) for Mouse hepatitis virus testing^27, 28^.

### Mouse lung endothelial cell isolation

The primary mouse lung endothelial cell (MLEC) was isolated from a 10-week-old, female, Balb/c mice and were cultured as described elsewhere^29^. Briefly, freshly isolated mouse lung were minced using autoclaved scissors, digested by collagenase I, and filtered through a 70-μm cell strainer. The cell suspension was incubated anti-rat Dynabeads (Thermo Fisher Scientific) conjugated with anti-mouse CD31 antibody (BD Biosciences, NJ, USA, 557355). Pooled cells were seeded in a 12-well-plate pre-coated with 0.1% gelatin. Upon reaching confluence, the cells were trypsinized and then incubated with anti-mouse ICAM-2 antibody (BD Biosciences, NJ, USA, 553326) conjugated Dynabeads. Pooled cells were seeded in 12-well-plate pre-coated with 0.1% gelatin. Harvested cells were assessed using tube formation assay, 1,1’-dioctadecyl-3,3,3’,3’-tetramethyl-indocarbocyanine perchlorate Low Density Lipoprotein (DiI-Ac-LDL) uptake, and CD31 gene expression (Figs. S4A - S4C).

### Tube formation assay

Tube formation was performed as described previously^30^. Briefly, 1 × 10^5^ JuB2 or MLEC suspended in 24-hour JuB2 conditioned medium were seeded in a 24-well plate pre-coated with Corning^®^ Matrigel^®^ Basement Membrane Matrix (Corning Inc. NY, USA). Cells were observed at 0, 2, 4, and 8 hours after seeding for tube formation.

### Dil-Ac-LDL uptake assay

Dil-Ac-LDL uptake assay was performed with Dil-Ac-LDL staining kit (Cell Applications, Inc.) according to the manufacturer’s instructions. Briefly, 6 × 10^5^ of isolated MLEC were cultured in a 4-well chamber slide (Nunc Lab-Tek Chamber Slide System, Thermo Fisher Scientific) precoated with Extracellular Matrix Attachment Solution provided in the kit. MLEC were allowed to grow until 95% confluency. Culture medium from each chamber was removed and cells were cultured in 100 µL of culture medium supplemented with 10 µg/mL of Dil-Ac-LDL. Cells were incubated for 4 hours, washed with wash buffer, and then mounted with DAPI-containing mounting medium and covered with 22 × 50 mm coverslip. Cells were examined under a confocal microscope (LSM700, Carl Zwiss, Oberkochen, German).

### Mice

All mouse experiments were performed under the AAALAC guidelines in Hokkaido University (protocol number:20-0083). Six-week-old male and female Balb/c mice purchased from Japan SLC, Inc. (Shizuoka, Japan) were used as breeders for tumor transplantation experiments. Mice were kept in a temperature-controlled specific-pathogen-free facility on a 12 hr light/dark cycle. Animals in all experimental groups were examined at least twice weekly for tumorigenesis.

### Tumor transplantation studies

ISOS-1, LEII, and UV♀2 cell lines were cultured in 15 cm dishes accordingly. Mice were randomly assigned to each group. 2 × 10^6^ ISOS-1, LEII, or UV♀2 cells were resuspended in Corning^®^ Matrigel^®^ Basement Membrane Matrix (Corning Inc. NY, USA) and inoculated subcutaneously in mice anesthetized with 3% isoflurane. Tumor sizes were measured twice weekly one week after inoculation. Mice were euthanized with CO_2_ when tumors reached 1,500 mm^3^ in volume or when mice exhibited abnormal behavior. Tumors were fixed in 10% neutral buffered formalin and processed for routine histological examination.

### Cell viability analysis

Cell viability after doxorubicin or GSK-J4 treatment was measured with Cell Counting Kit-8 (Dojindo Molecular Technologies, Inc., Kumamoto, Japan) according to the manufacturer’s instructions. The absorbance at 450 nm was measured with NanoDrop™ 2000 (Thermo Fisher Scientific). Determination of IC_50_ were performed using KyPlot 6.0 software (KyensLab, Inc., Tokyo, Japan). Experiments were performed at least three times with triplicates.

### Western blotting

Western blotting was performed as described previously^7^. The antibodies used in this study are as follows: anti-KDM1A antibody (1:1000; Cell Signaling Technology, MA, USA, 2139S), anti-KDM2A antibody (1:1000; Abcam, Cambridge, UK, Ab191387), anti-KDM2B antibody (1:1000; Santa Cruz Biotechnology, Inc., TX, USA, sc-293279), anti Actin antibody (1:10000; Sigma Aldrich, MO, MAB1501), anti-c-FOS antibody (1:1000; Santa Cruz Biotechnology, Inc., sc-166940), anti-γH2A.X antibody (1:1000; Bethyl Laboratories, Inc, TX, USA. A300-081A-T), anti-p-ERK1/2 antibody (1:1000; Cell Signaling Technology, 4370S), anti-ERK1/2 antibody (1:1000; Cell Signaling Technology, 4695S), anti-H2AK119Ub1 antibody (1:2000; Cell Signaling Technology-8240S), anti-H3 antibody (1:3000; MAB Institute, Inc. Yokohama, Japan, MABI0001-20), and CD204 (1:125; Medicinal Chemistry Pharmaceutical Co., Ltd., Sapporo, Japan, KT022). All primary antibodies were diluted in Can Get Signal Solution^®^ 1 (TOYOBO, Osaka, Japan). Membranes were washed with Tris-Buffered Saline with 0.1% Tween^®^ 20 (TBS-T) three times for 5 mins each time before incubating with ECL Mouse IgG HRP-linked whole antibody (Cytiva, MA, USA, #NA934) or ECL Rabbit IgG HRP-linked whole antibody (Cytiva, #NA931) diluted in Can Get Signal Solution 2 (TOYOBO, Osaka, Japan). Signal development was performed using Immobilon^®^ Western Chemiluminescent HRP substrate (Merck Millipore, NJ, USA). ImageQuant LAS 4000 mini luminescent image analyzer (GE Healthcare) was used to visualize chemiluminescent signals and the ImageJ software was used to process captured data^31^.

### Quantitative RT-PCR (qRT-PCR)

qRT-PCR was performed as described previously^7^. The list of primers used in this study is listed in Supplementary table. cDNA of mouse mesenchymal stem cells (mMSC), donated by Dr. Yusuke Komatsu (Hokkaido University), were used as a negative control for *CD31* expression.

### Flow cytometry analysis

Flow cytometry for cell cycle was performed as described previously^7^. Briefly, 2 × 10^5^ HSA cells were harvested for each replicate and unstained control. Samples were fixed with 70% ethanol and incubated with propidium iodide (PI) in the dark for 30 mins at 37°C. Unstained cells were incubated with PBS for 30 mins at 37°C. Cell cycle was analyzed in BD FACSVerse™ flow cytometer (BD Biosciences, NJ, USA). Results were analyzed with FCS Express 4 software (De Novo Software, CA, USA). Experiments were performed at least three times with triplicates.

For PD-L1 expression analysis, HSA cells were cultured in 6-well plates until 90% confluency. Cells were washed with 1.34 mM EDTA in PBS twice and then detached by adding 1 mL of 1.34 mM EDTA solution and incubating at room temperature (RT) for 10-15 mins. Cells were counted and 2 × 10^5^ cells were used for each replicate. Cells were blocked with 10% goat serum in PBS with sodium azide at RT for 15 mins before incubating with anti PD-L1 antibody (1:100; clone 6C11-3A11) or isotype rat IgG2a control (1:50; BD Biosciences) at RT for 30 mins^21^. Cells were washed with 1% BSA in PBS twice before incubating with secondary anti-rat IgG antibody conjugated with APC (1:500; Southern Biotech, AL, USA, Catalog no. 3010-11L). Cells were analyzed in BD FACSVerse™ flow cytometer (BD Biosciences, NJ, USA). Results were analyzed with FCS Express 4 software (De Novo Software, CA, USA). Experiments were performed at least three times with triplicates.

### RAW264.7 polarization

RAW264.7 cells were polarized to M1 or M2 macrophages as described by a previous study^32^. 1 × 10^6^ RAW264.7 cells were seeded in 10 cm dishes and were stimulated with LPS (100 ng/ml) and IFN-γ (20 ng/ml) or TGF-β (10 ng/ml) for 24 hours for their polarization towards the M1 or M2 subtype, respectively. RAW264.7 cells were also incubated in 48 hour-conditioned medium from ISOS-1 cell culture filtered with a 0.45 µm sterile filter. RAW264.7 cells cultured under normal DMEM medium were considered as M0 or unpolarized macrophages.

### Transwell migration assay

Migration assay for RAW264.7 cells was performed as described previously with minor changes^33^. Briefly, 2.5 × 10^5^ RAW264.7 cells in 1 mL normal medium were added to ThinCerts cell culture inserts (Greiner Bio-One, Kremsmünster, Austria) in 6 well plates while 2.5 mL of 48 hours conditioned medium from MLEC or ISOS-1 cells were added to the lower wells. After 24 hours, cells in the upper wells were removed with a cotton swab, filters were fixed with 4% paraformaldehyde in PBS and stained with 0.1% crystal violet solution in DW. Cell migration was assessed by counting the number of migrated cells in five fields per well at 40× magnification.

### Histopathological analysis and Immunohistochemistry

Tissue samples were fixed for at least 48 hours in 10% neutral buffered formalin, processed routinely, embedded in paraffin, and sectioned 3 μm thick, mounted on glass slides, and stained with hematoxylin and eosin (HE). All tumors were examined for KDM2B expression and presence of lymphocytes and macrophages using immunohistochemistry (IHC). IHC was performed using antibodies to Iba1 (1:2000, Fujifilm Wako, Osaka, Japan, 019-19741), CD3 (1:1000; Agilent Technologies, CA, USA, IR503), KDM2B (1:50; Santa Cruz Biotechnology, Inc.,sc-293279), CD31 (1:250; Abcam, JC/70A), PD-L1 (1:100; clone 6C11-3A11), and CD204 (1:800; Medicinal Chemistry Pharmaceutical Co., Ltd., Sapporo, Japan, KT022) as described previously^7^. Nano Zoomer 2.0-RS (Hamamatsu Photonics, Hamamatsu, Japan) was used to scan histological slides, which were then processed in QuPath ver 0.2.133^34^. Scanned slides were opened in QuPath as Brightfield (H-DAB), and the Estimate Stain Vectors feature was used to automatically adjust the staining colors. Normal endothelial and tumor cells were detected using the Cell Detection function. The Create Detection Classifiers function was used to label cells based on their morphologies and locations, allowing the QuPath software to correctly classify each cell type. The collected data was exported and used for further analysis.

### Immunofluorescence assay

Tissue samples were fixed, processed, and incubated with primary antibodies for Iba1 and PD-L1 as described above. Tissue samples were washed with PBS three times for 5 minutes each before adding a secondary goat anti-rabbit IgG H & L (1:1000, Alexa Fluor® 555, Abcam) and Goat Anti-Rat IgG H & L (1:1000, Alexa Fluor® 488, Abcam) in 5% skim milk for 1 hour at RT. Sections were washed with PBS three times for 5 mins each, mounted with DAPI-containing mounting medium, and covered with 24 × 32 mm coverslip. Tissues were examined under a confocal microscope (LSM700, Carl Zwiss, Oberkochen, German).

### Statistical analysis

Statistical analyses were performed with Microsoft Excel and R software (version 3.6.3). Student’s *t*-test was used to analyze the difference between two groups while Tukey’s test was used to analyze differences between multiple groups. *P-*values less than 0.05 were considered statistically significant.

## Data availability

The datasets generated during and/or analyzed during the current study are available from the corresponding author on reasonable request.

## Acknowledgments

We would like to extend our sincerest gratitude to Dr. Kazuhiro Kimura (Hokkaido University) and Dr. Hiroki Sakai (Gifu University) for providing LEII cell line and canine hemangiosarcoma cell lines, respectively. We are grateful to the members of the Laboratory of Comparative Pathology, Faculty of Veterinary Medicine, Hokkaido University for their invaluable support during the conduct of this study. This research was supported by the KAKENHI Grant-in-Aid for Young Scientist (KA, 20K15654) provided by Japan Society for the Promotion of Science, and by a research grant provided by the Kuribayashi Foundation (KA, No. 2-2)

## Author contributions

**K. G**. Conceptualization, Methodology, Investigation, Formal Analysis, Data Curation, Writing-Original Draft, Writing-Review & Editing, **K. A**. Conceptualization, Writing-Review & Editing, Supervision, Funding acquisition. **N. M**. and **S. K**. Methodology and supervision. **A. K**. Writing-Review & Editing. **T.K**. Supervision.

## Figure legends

**Table.** PD-L1 expression in tumor and immune cells in canine HSA.

| Patient No. | Sex | Age (years) | Breed | PD-L1 intensity (tumor cells) | PD-L1 intensity (macrophages) | PD-L1 intensity (lymphocytes) |
| --- | --- | --- | --- | --- | --- | --- |
| 1 | Male (castrated) | 7 | Miniature Dachshund | - | +++ | - |
| 2 | Female (spayed) | 7 | Golden Retriever | + | - | - |
| 3 | Male | 14 | Miniature Schnauzer | - | - | - |
| 4 | Female (spayed) | 10 | Mixed | + | +++ | - |
| 5 | Male (castrated) | 12 | Beagle | + | - | - |
| 6 | Male | 8 | Labrador Retriever | ++ | +++ | - |
| 7 | Female (spayed) | 10 | Miniature Dachshund | ++ | +++ | - |
| 8 | Female (spayed) | 14 | Miniature Dachshund | - | +++ | - |
| 9 | Female (spayed) | 14 | Miniature Schnauzer | - | - | - |
| 10 | Male (castrated) | 12 | Miniature Schnauzer | + | +++ | - |
| 11 | Male | 13 | French Bulldog | - | - | - |
| 12 | Male (castrated) | 14 | Miniature Dachshund | - | +++ | - |
| 13 | Female (spayed) | 12 | Labrador Retriever | - | +++ | - |
| 14 | Male (castrated) | 14 | Lhasa Apso | + | - | - |
| 15 | Female (spayed) | 12 | Beagle | - | +++ | - |
| 16 | Female (spayed) | 14 | Miniature Dachshund | + | +++ | - |
| 17 | Male (castrated) | 14 | Miniature Dachshund | ++ | +++ | - |
| 18 | Female | 8 | Flat-coated Retriever | - | +++ | - |
| 19 | Female (spayed) | 11 | Scottish Terrier | ++ | +++ | - |
| 20 | Female | 5 | Golden Retriever | ++ | +++ | - |
| 21 | Male | 5 | Beagle | ++ | +++ | - |
| 22 | Female | 9 | Golden Retriever | + | - | - |
| 23 | Male | 10 | Miniature Schnauzer | ++ | +++ | - |
| 24 | Female | 12 | Mixed | - | - | - |
| 25 | Male | 9 | Great Pyrenees | - | +++ | - |
| 26 | Male | 7 | Jack Russel Terrier | - | - | - |
| 27 | Male (castrated) | 8 | Labrador Retriever | - | +++ | - |
| 28 | Male (castrated) | 10 | Maltese | + | +++ | - |

