## Supplementary files for "Hemangiosarcoma cells induce M2 polarization and PD-L1 expression in macrophages"

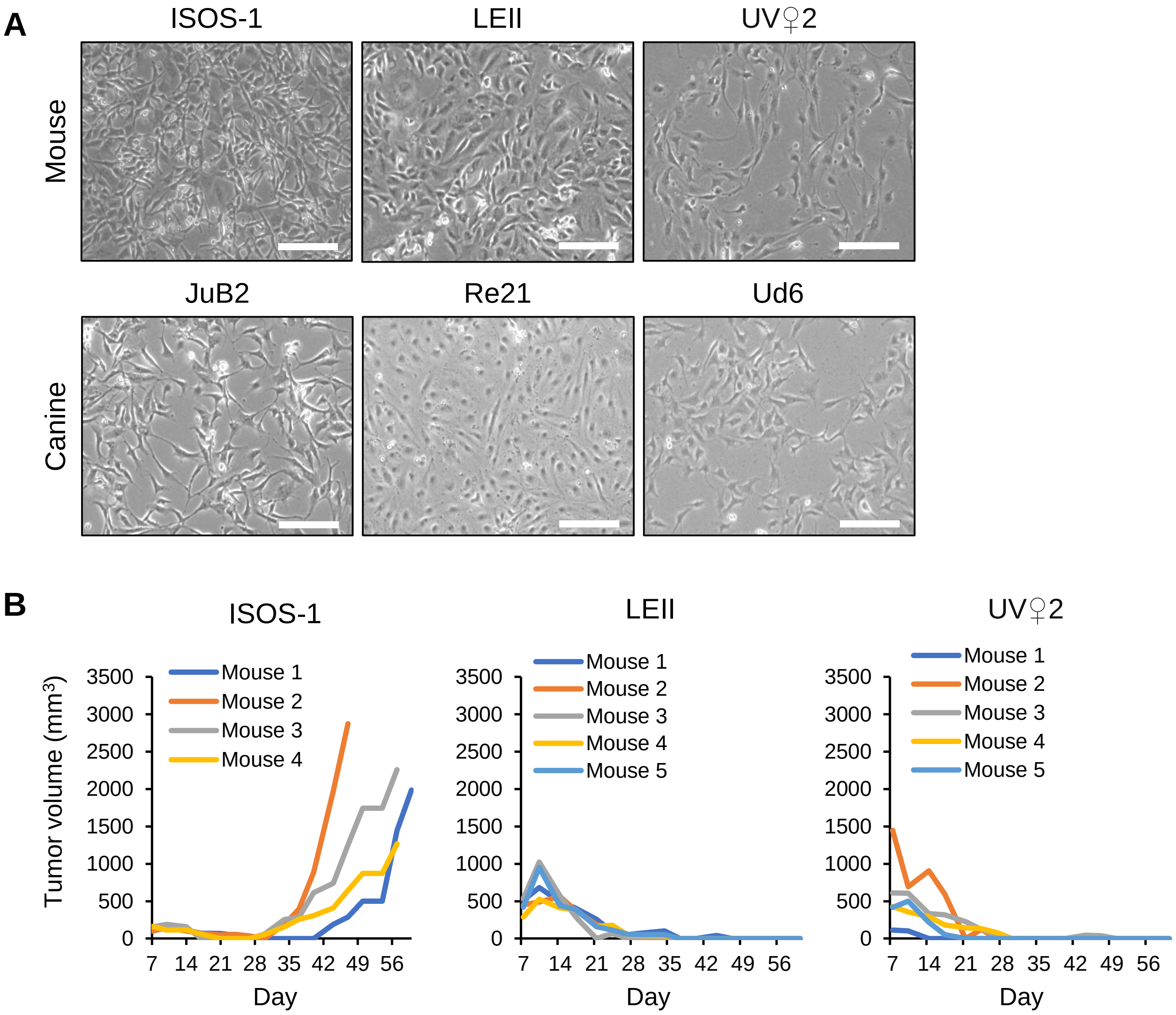


**Supplementary Figure 1. Morphologies of mouse and canine HSA cell lines and tumor growth curves of mouse HSA cell lines.**

**A,** Phase-contrast images of mouse HSA cells (ISOS-1, UV♀2), mouse immortalized endothelial cells (LEII), and canine HSA cell lines (JuB2, Re12, Ud6). **B,** Tumor growth curves of ISOS-1, LEII and UV♀2 in Balb/c mice. Bars = 125μm.

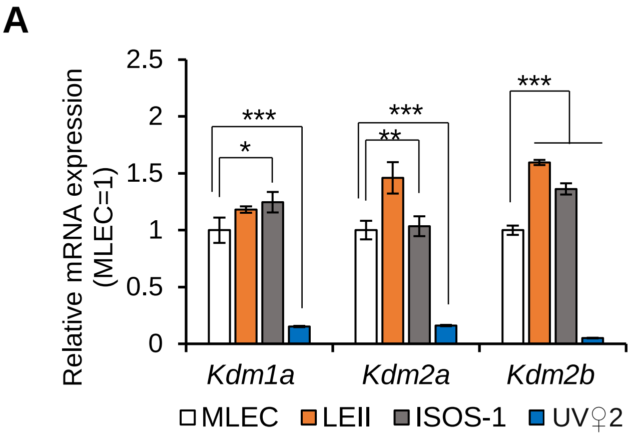


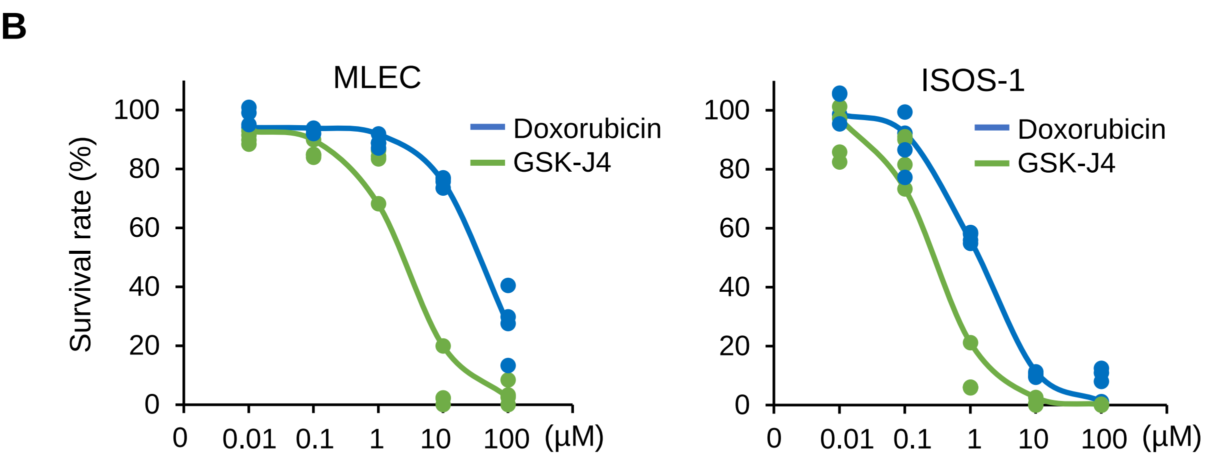


| Drug | $IC_{50}$ |
|---|---|
| Doxorubicin | 77.39 μM |
| GSK-J4 | 5.06 μM |

| Drug3 | $IC_{50}$ |
|---|---|
| Doxorubicin | 2.56 μM |
| GSK-J4 | 0.53 μM |

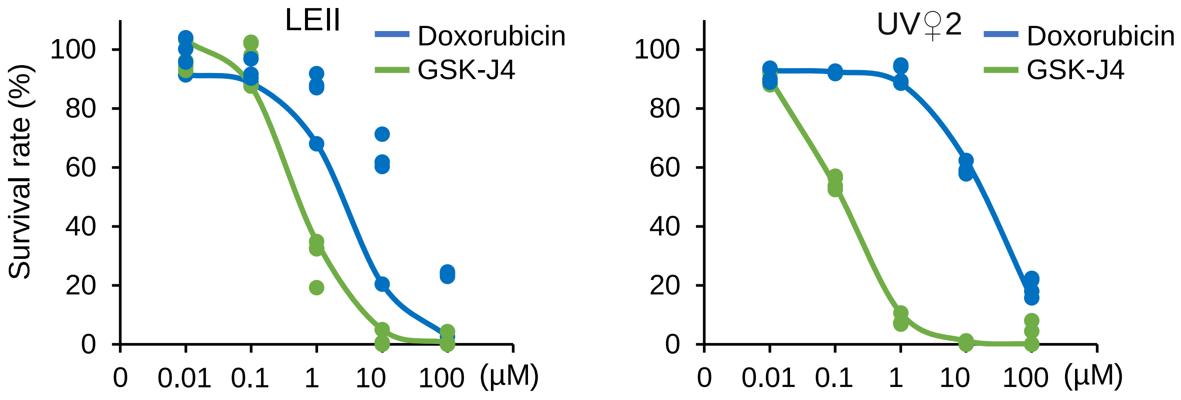


| Drug | $IC_{50}$ |
|---|---|
| Doxorubicin | 5.25 μM |
| GSK-J4 | 1.04 μM |

| Drug | $IC_{50}$ |
|---|---|
| Doxorubicin | 38.05 μM |
| GSK-J4 | 0.24 μM |

**Supplementary Figure 2. Molecular similarity of mouse HSA cell lines with canine HSA cell lines.** **A,** Relative gene expressions of *Kdm1a, Kdm2a,* and *Kdm2b* in mouse ISOS-1, LEII and UV♀2.. **B**, Survival rates and $IC_{50}$ values of doxorubicin- or GSK-J4-treated MLEC, ISOS-1, LEII and UV♀2. Data are presented as mean values ± s.d. $^{**}$ $P< 0.01^{***}$ $P< 0.001$, Tukey's test

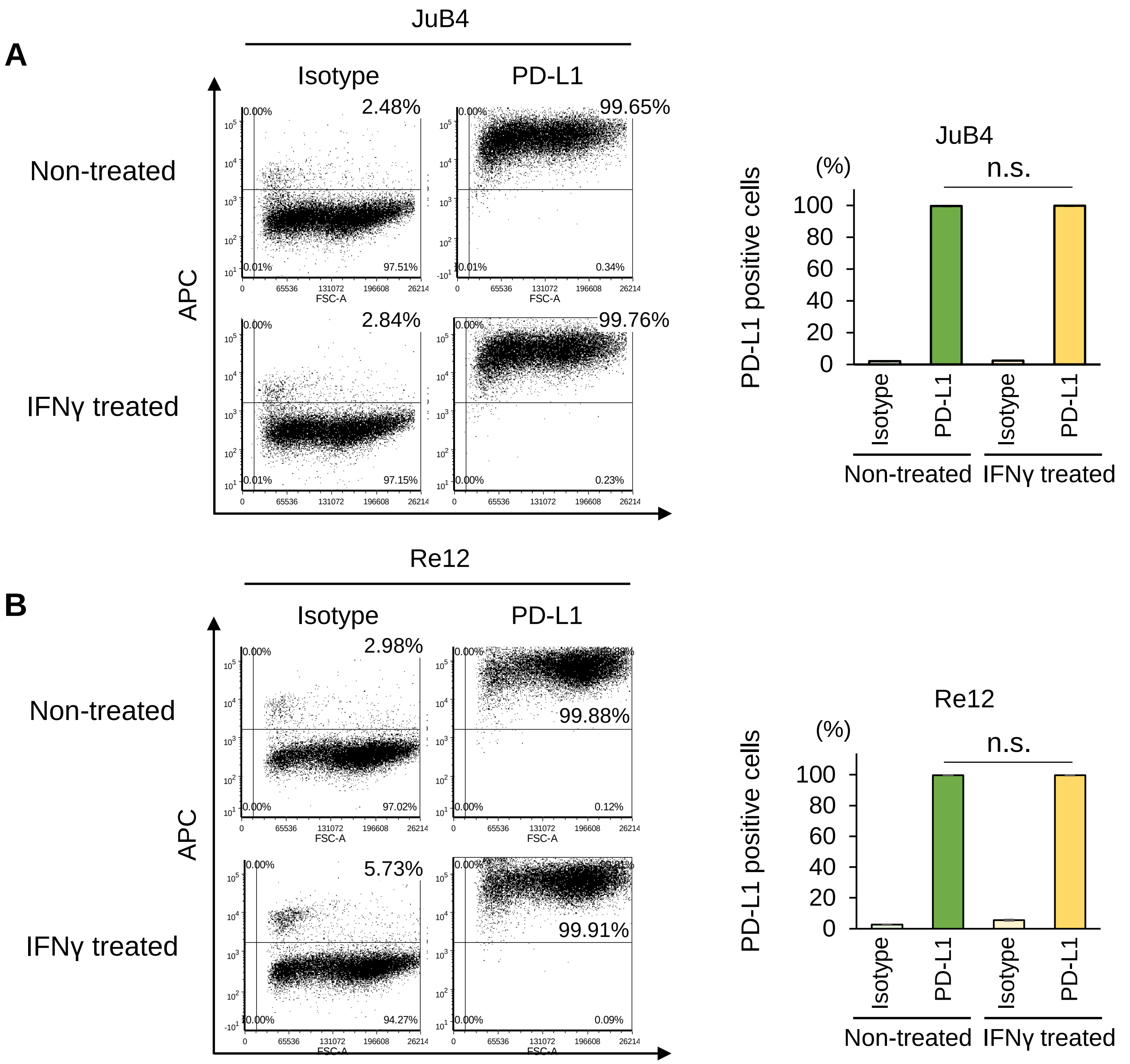


**Supplementary Figure 3. PD-L1 is constitutively expressed in canine HSA cell lines.**
**A and B,** (Left) Representative images of flow cytometry for PD-L1 in JuB4 (A) and Re12 (B) cell lines with/without IFNγ. (Right) Quantitative analysis of the flow cytomery data. Data are presented as mean values ± s.d.

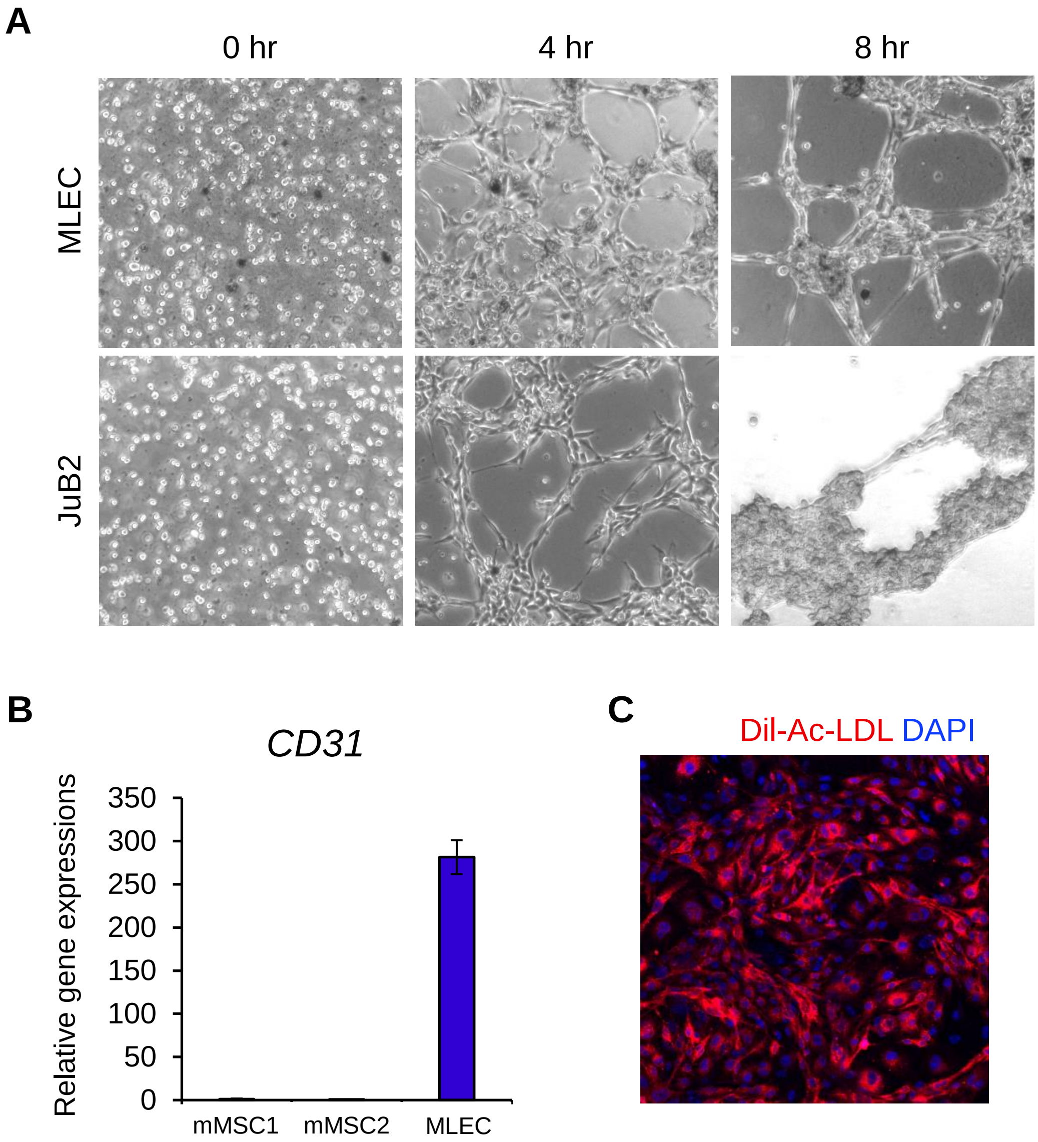


**Supplementary Figure 4. Verification and characterization of isolated MLEC.**
**A,** Phase-contrast images of tube formation assay for MLEC and JuB2. **B,** Relative gene expression levels of *CD31* in mouse mesenchymal stem cells (mMSC) and MLEC. **C,** Dil-Ac-LDL uptake assay in isolated MLEC. Data are presented as mean values ± s.d.

| Species | Target | Sequence |
|---|---|---|
| Mouse | Kdm1a (F) | TTCCCAGACATCATCAGTGG |
| | Kdm1a (R) | CCAGCCATAACTGCAATGTG |
| | Kdm2a (F) | ACTGCATAACCAACCGATCC |
| | Kdm2a (R) | CCTTCCTCATCACCATTTCC |
| | Kdm2b (F) | TCCTGCATAGCTTCAACGTG |
| | Kdm2b (R) | TAACGGAACTTGGGCTGAAC |
| | Tbp (F) | AACAGCCTTCCACCTTATGC |
| | Tbp (R) | AAGATGGGAATTCCAGGAGTC |
| | CD31 (F) | CCAAAGCCAGTAGCATCATGGTC |
| | CD31 (R) | GGATGGTGAAGTTGGCTACAGG |
| | IL-6 (F) | TACCACTTCACAAGTCGGAGGC |
| | IL-6 (R) | CTGCAAGTGCATCATCGTTGTTC |
| | Nos2 (F) | GAGACAGGGAAGTCTGAAGCAC |
| | Nos2 (R) | CCAGCAGTAGTTGCTCCTCTTC |
| | Tnfα (F) | GGTGCCTATGTCTCAGCCTCTT |
| | Tnfα (R) | GCCATAGAACTGATGAGAGGGAG |
| | Havcr2 (F) | ACAGACACTGGTGACCCTCCAT |
| | Havcr2 (R) | CAGCAGAGACTCCCACTCCAAT |
| | CD163 (F) | GGCTAGACGAAGTCATCTGCAC |
| | CD163 (R) | CTTCGTTGGTCAGCCTCAGAGA |
| | PD-L1 (F) | TGCGGACTACAAGCGAATCACG |
| | PD-L1 (R) | CTCAGCTTCTGGATAACCCTCG |

**Supplementary table**. Primers used in this study.
